# Ultrasensitive tuning of cytoplasmic viscosity via active noise

**DOI:** 10.1101/2024.12.05.626947

**Authors:** Ziming Zhao, Jie Lin

## Abstract

The mechanical properties of cytoplasm are crucial for cellular functions. While active processes significantly alter cytoplasmic viscoelasticity, the physical mechanisms remain elusive. Here, we model the cytoplasm as a colloidal suspension subject to passive and active noise, coarse-grained as an effective temperature. We show that a jammed cytoplasm transitions from a solid to a liquid phase at a critical effective temperature. Intriguingly, the simulated complex shear modulus at the critical state exhibits a 1*/*2 power-law scaling with frequency and quantitatively matches the experimental data of live cells without fitting parameters, given that the cytoplasmic volume fraction is slightly above the jamming transition. We further reveal the biological significance for the cytoplasm to be slightly above jamming: it is a regime in which the viscosity is ultrasensitive to changes in the effective temperature. Our results suggest that cells actively regulate their volume fractions in the sensitive regime in which they can tune cytoplasmic viscosity efficiently via active processes.

## Introduction

The mechanical properties of cytoplasm are crucial determinants of its physiological functions in diverse biological contexts [1–7]. Understanding the factors that regulate the viscoelasticity of cytoplasm is thus paramount for comprehending cell behaviors and functions. Experiments have suggested that cytoplasm is a crowded suspension of biomolecules resembling colloidal systems of deformable particles [8–10]. A critical feature that makes cytoplasm distinct from passive systems is the active, energy-consuming processes inherent to live cells. Experiments based on microrheology have measured the viscoelastic properties of cytoplasm for mammalian cells [11–13]. Interestingly, the storage modulus of live cells is significantly lower than that of ATP-depleted cells at low frequency: active processes fluidize the cytoplasm from a solid-like state, also observed in bacteria [7]. Furthermore, both the storage and loss moduli exhibit a power-law scaling with frequency and are approximately equal over a large range of frequency: *G*^*′*^ (*ω*) ≈ *G*^′′^(*ω*) ∼ *ω*^1*/*2^ [11, 13, 14]. In this sense, the cytoplasm is in a critical state, neither solid nor liquid. However, the physical origin of the *ω*^1*/*2^ scaling is still unclear and controversy [11, 15]. Furthermore, the effective volume fraction of cytoplasm as a colloidal suspension is still unknown, and whether the volume fraction of cytoplasm is regulated to be an optimal value in live cells is even more elusive.

In this study, we study the influence of cytoplasmic density and noise generated by active intracellular processes on the viscoelastic properties of cytoplasm (Figure 1a). We model the cytoplasm as a suspension of soft, spherical particles (Figure 1b), incorporating passive and active noise, which we coarse-grain together as an effective temperature. We compute the complex shear modulus and viscosity as a function of the effective temperature and volume fraction, *ϕ* (the relative volume occupied by particles). This simple model has been intensively studied: at zero temperature, it exhibits a jamming transition from the liquid phase to the solid phase at *ϕ*_*J*_ *≈* 0.64 for monodisperse systems in three dimensions [16–18]. We show that systems above jamming with *ϕ > ϕ*_*J*_ can be fluidized by noise and exhibits a transition from the solid phase to the liquid phase at the critical effective temperature *T*_eff,c_. At *T*_eff,c_, the shear stress upon a constant strain relaxes as a power-law function of time, *σ* ∼ *t*^−1*/*2^, in analogy to the critical state at *ϕ*_*J*_ and *T*_eff_ = 0 [19, 20]. Surprisingly, the simulated complex shear modulus at *T*_eff,c_ quantitatively matches the experimental data of mammalian cells without any fitting parameters as long as we take a volume fraction slightly above *ϕ*_*J*_. Interestingly, we find that the active part dominates the effective temperature, in concert with the non-equilibrium nature of cytoplasm [12, 13, 21, 22].

**FIG. 1.**
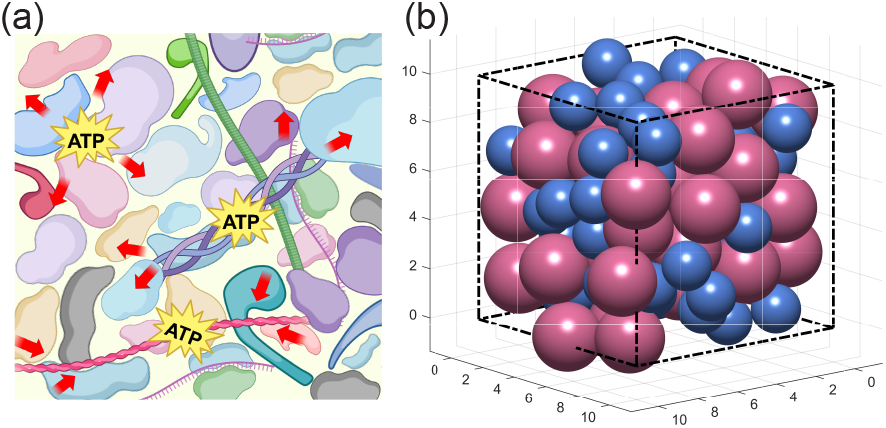
(a) A schematic of cytoplasm with active noise generated by ATP-consuming processes, such as transportation processes along the cytoskeleton. (b) We model the cytoplasm as a colloidal suspension of bidisperse particles subject to passive and active noises.

To understand whether there are any biological functions for cells to be slightly above jamming, we compute the viscosity, the ratio between the stress and the shear rate in the long-time limit, as a function of volume fraction and effective temperature. Intriguingly, we reveal a sensitive regime of volume fraction slightly above *ϕ*_*J*_ in which the viscosity is ultrasensitive to the effective temperature. We propose that cells actively regulate their volume fractions to be in the sensitive regime to efficiently tune their cytoplasmic viscosity via a slight change in the amount of intracellular active processes to achieve certain biological functions. Further, the active noise also makes the cytoplasmic viscosity less sensitive to the change in volume fraction, in agreement with previous experiments of live cells [8, 10] and passive colloidal systems [9].

### A particle-based model of cytoplasm

We model the cytoplasm as a colloidal suspension of soft spherical particles whose dynamics follow the overdamped Langevin equation:

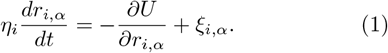

Here, *i* = 1, 2, …, *N* where *N* is the number of particles, and *α* = *x, y, z* the directions in the Cartesian coordinate. *η*_*i*_ is the friction coefficient, which is related to the viscosity of the background solvent through the Stokes’ relationship, *η*_*i*_ = 6*πνa*_*i*_. Here, *a*_*i*_ is the radius of particle *i*, and *ν* is the solvent viscosity. We use a 50-50 mixture of particles with a size ratio of 1.4 to prevent crystallization [16, 23]. Particles interact through the “one-side spring” model, *U* = Σ_*i>j*_*κ*(*a*_*i*_ +*a*_*j*_ − *r*_*ij*_)^2^Θ(*a*_*i*_ +*a*_*j*_ − *r*_*ij*_)*/*2, where *r*_*ij*_ = |**r**_*i*_− **r**_*j*_ |, *κ* is the spring constant, and Θ is the Heaviside function. The volume fraction quantifies the crowdedness of the cytoplasm, defined as 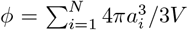, where *V* is the system volume. The fluctuating force *ξ*_*i*,*α*_ is modeled as Gaussian noise satisfying the Fluctuation-Dissipation (FD) theorem [24]

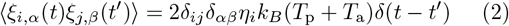

Here, *k* is the Boltzmann constant, and *T*_p_ is the passive temperature, i.e., thermodynamic temperature. We introduce *T*_a_ as the active temperature, an effective variable coarse-graining the effects of active intracellular processes, e.g., ATP-consuming processes (Figure 1a). Here, *δ*_*ij*_ and *δ*_*αβ*_ are Kronecker delta function and *δ*(*t*− *t*^′^) is Dirac delta function. We include the details of numerical simulations in the Supplementary Material.

We remark that the strength of thermal noise is proportional to the particle radius because the friction coefficient is proportional to the radius in Eq. (2). One of us has recently shown that the strength of active noise may be proportional to the cube of particle radius [25]. Since the particle sizes are similar in most parts of this work, we simplify the model by combing the thermal noise and active noise together and define *T*_eff_ = *T*_p_ + *T*_a_ as the effective temperature incorporating both thermal and active fluctuations. We non-dimensionalize the system such that the length unit is the radius of the small particles *a*_0_, the time unit *t*_0_ = 6*πνa*_0_*/κ* and the energy unit 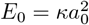 In the following, all variables are dimensionless unless otherwise mentioned.

### Noise fluidizes jammed cytoplasm

In the following, we study the viscoelastic response of the cytoplasm model. In this section, we do not specify the contribution of thermal and active noises to the effective temperature. We later show that the experimental data of live mammalian cells suggest that the active part dominates the effective temperature [11]. We first let the system relax for a time *t* = 5 × 10^3^ and then apply a small strain to the system through the Lees-Edwards boundary condition: particles initially at (*x, y*) are displaced to (*x, y* +*ϵx*) [26]. We compute the stress as *σ*(*t*) = Σ_*i>j*_*F*_*ij*,*x*_(*y*_*i*_ − *y*_*j*_)*/L*^3^, where *F*_*ij*,*x*_ represents the elastic force of particle *i* on *j* and monitor the stress relaxation as a function of time, *σ*(*t*), from which we compute the viscosity *η* and complex shear modulus *G*^*^ as

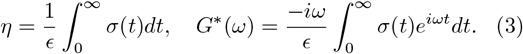

Our results agree with previous works for the zero-temperature case. The bidisperse system exhibits the jamming transition at a critical volume fraction *ϕ*_*J*_ ≈0.648 [16]. Below jamming (*ϕ < ϕ*_*J*_), the shear stress relaxes to zero; above jamming (*ϕ > ϕ*_*J*_), the shear stress relaxes to a finite value *σ*_∞_, which represents the unrelaxable part of stress; at the jamming point (*ϕ* = *ϕ*_*J*_), the shear stress decays *σ*(*t*) ∼ *t*^−1*/*2^ (Figure S1) [20]. We note that the 1*/*2 power-law decay comes from the abundance of soft modes of the Hessian matrix *H*_*iα*,*jβ*_ = *∂*^2^*U/∂r*_*iα*_*∂r*_*jβ*_ [27]; in particular, the distribution of relaxation rates over eigenmodes satisfies *D*(*s*) ∼ *s*^−1*/*2^ for *s <* 1 at the jamming point [19, 28]. Therefore, one can approximate the stress as

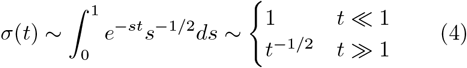

where we assume that the contribution of each eigenmode to stress is comparable. For systems above jamming, the − 1*/*2 temporal scaling of stress crossovers to a plateau in the long time limit [20] (Figure S2).

We next study jammed systems with *ϕ > ϕ*_*J*_ to demonstrate the fluidization effects of noise. We fix *ϕ* and tune the effective temperature *T*_eff_ over two decades and calculate the normalized relaxation curve *σ*(*t*)*/σ*(0) (Figure 2a). Interestingly, we also find a transition from a solid phase to a liquid phase as *T*_eff_ increases. At the critical temperature *T*_eff,c_, the stress also decays as *σ*(*t*) ∼ *t*^−1*/*2^: the stress exhibits the same power-law decay at *T*_eff,c_ as at the jamming point without noise. Similar results are obtained if we fix *T*_eff_ and tune *ϕ* (Figure S3). Given the interaction form between particles, the elastic energy scales with the distance to the jamming point as *U* ∼ (*ϕ* − *ϕ*_*J*_)^2^ [29]. The simplest scenario is that the average energy barrier for escaping a stable state has the same scaling as the energy itself [30], leading to the following scaling between the critical temperature *T*_eff,c_ and volume fraction *ϕ*

**FIG. 2.**
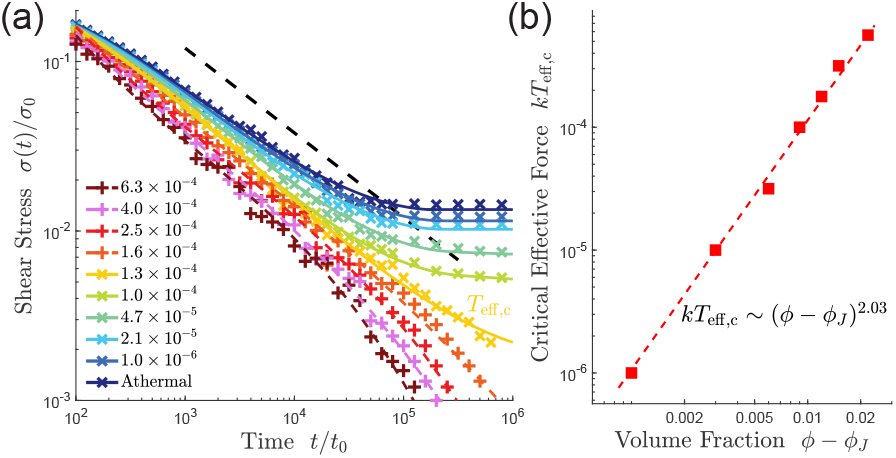
(a) A jammed cytoplasm with *ϕ* = 0.660 is fluidized by noise such that the stress relaxes to zero at a high enough effective temperature. The critical effective temperature *T*_eff,c_ ≈1.3 × 10^−4^ in this case. The colorful dashed lines represent a third-order polynomial fitting of the simulated data. The black dashed line has a slope of −0.5 in this log-log plot. (b) The scaling between the critical temperature *T*_eff,c_ and *ϕ* − *ϕ*_*J*_ where *ϕ*_*J*_ is the volume fraction at the jamming transition at zero temperature.

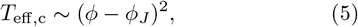

confirmed by our simulations (Figure 2b).

### Live cells have jammed cytoplasm and resemble critical jamming system

Recent experiments of active microrheology have measured the viscoelasticity properties of live cells in which one inferred the complex shear modulus by applying a force on a probe particle in live cells and measuring its displacement [11–13]. Interestingly, Ebata et al. found that the storage modulus of ATP-depleted cells is significantly higher than that of live cells at low frequency (*ω <* 10 Hz): the cytoplasm is solid-like if cellular activities are suppressed and fluidized by active noise. Similar phenomena have also been observed in bacteria [7]. More interestingly, both the storage and loss moduli of live cells exhibit approximately power-law scaling with frequency, *G*^*′*^ (*ω*) ∼ *ω*^1*/*2^ and *G*^*′′*^ (*ω*) ∼ *ω*^1*/*2^ at low frequency. We remark that the *ω*^1*/*2^ scaling of shear moduli comes from the stress scaling *σ*(*t*) ∼ *t*^−1*/*2^ after a constant strain according to Eq. (3), which suggests that the volume fraction of cytoplasm is likely above jamming, but the system still resembles a critical system due to active noise (Figure 2a).

To verify our hypothesis, we compare the simulated complex moduli at the critical effective temperature with the experimental data. Surprisingly, the simulated data matches the experimental data well with *ϕ* = 0.660 and *T*_eff,c_ ≈1.3 × 10^−4^ for both the storage modulus and the loss modulus (Figure 3a, b). We also infer *T*_p_ ≈2.5 × 10^−5^ from the fitting of ATP-depleted cells given the same *ϕ*. In particular, our results suggest that the volume fraction of cytoplasm should be slightly above the jamming point. If the volume fraction is too high, an unreasonably high temperature must match the experimental data (Figure S4). On the other hand, if the volume fraction is below jamming, increasing effective temperature makes the system more solid-like, in constant to experiments, which we discuss in the next section. Our analysis shows that the active part in live cells dominates the effective temperature, and this conclusion is independent of *ϕ* as long as it is in the sensitive regime (Figure S5). In the next section, we show that a volume fraction slightly above jamming is a sensitive regime in which the viscosity is ultra-sensitive to the effective temperature.

**FIG. 3.**
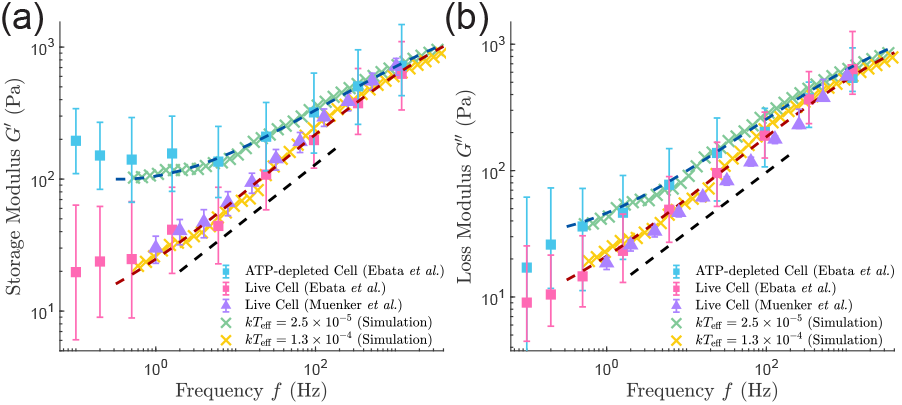
The simulated complex moduli with *ϕ* = 0.660 at the critical temperature *T*_eff,c_ match the experimental data well, including the storage modulus (a) and the loss modulus (b). Given the same *ϕ*, we infer the passive temperature *kT*_p_ ≈2.5 × 10^−4^ from the fitting of ATP-depleted cells. The experimental data are from Ref. [11] and Ref. [13]. The colorful dashed lines represent a third-order polynomial fitting of the simulated data. The black dashed lines have a slope of 0.5 in the log-log plot.

### Ultra-sensitivity of cytoplasmic viscosity to active noise

We next seek to understand the biological function of active noise and why the volume fraction of cytoplasm is slightly above jamming. We compute the viscosity using Eq. (3) as a function of effective temperature and volume fraction (Figure 4a) [31]. In contrast to our results for jammed systems, for systems below jamming (*ϕ < ϕ*_*J*_), increasing the effective temperature enhances the viscosity (Figure 4b). This “counter-intuitive” observation results from the glass transition: thermal fluctuation increases viscosity for systems below jamming [32]. We propose that evolutionary selection keeps the cytoplasm above jamming; otherwise, cellular activities would make the cytoplasm more viscous instead of more liquid-like, which is presumably biologically unfavorable and also contradicts experiments [7, 11].

**FIG. 4.**
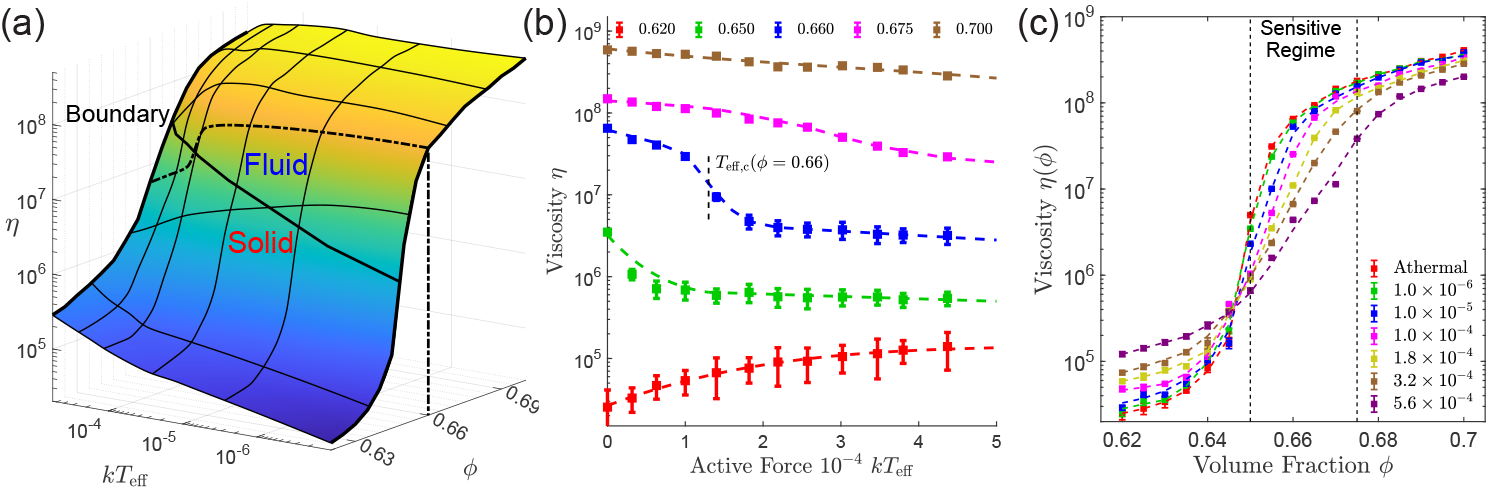
(a) Viscosity as a function of the volume fraction and effective temperature. The dash-dot line represents the viscosity at *ϕ* = 0.66. The solid line is the critical effective temperature *T*_eff,c_. (b) Viscosity vs. *T*_eff_ under different volume fractions. Below jamming, the viscosity increases with temperature. Well above jamming, the viscosity decreases mildly with temperature. Slightly above jamming, the viscosity exhibits a sharp decrease at the critical effective temperature (the vertical dashed line).(c) Viscosity vs. *ϕ* under different effective temperatures. We highlight the sensitive regime slightly above the jamming point *ϕ*_*J*_ ≈0.650. In this regime, the slope of growth decreases as effective temperature increases, indicating that a finite active noise also makes the viscosity less sensitive to the change in *ϕ*.

Meanwhile, a volume fraction well above jamming is not evolutionarily advantageous either since a very high effective temperature is required to tune the cytoplasmic viscosity in a limited range (Figure 4b). Intriguingly, we find a sensitive regime slightly above *ϕ*_*J*_ (Figure 4a), in which the viscosity exhibits a significant change near the critical effective temperature (Figure 4b, c). Taken together, we propose that cells adjust their cellular activities to lie at the border between the solid and liquid phases, allowing efficient cytoplasmic viscosity tuning, in agreement with the *ω*^1*/*2^ scaling of shear moduli in live cells [11, 13].

On the other hand, the cell volume can change upon osmotic or mechanical perturbations; the resulting change in volume fraction disturbs the viscosity and impacts cellular physiology globally. A finite active noise also makes the system less sensitive to the change in volume fraction in the sensitive regime (Figure 4c), in agreement with experiments of live cells [8, 10] and deformable colloidal suspensions [9]. Our results demonstrate that active processes enhance cellular robustness to overcome adverse environmental impacts on cell volume.

## Discussions

In this work, we model the cytoplasm as a suspension of repulsive particles and show that the cytoplasm of live cells may be slightly above jamming and poised to the critical state by active noise. The criticality leads to ultrasensitivity of cytoplasmic viscosity to active noise. Therefore, cells can effectively tune their viscosities and adapt to different environments, essential for biological processes such as tumor invasion and cell dormancy [33, 34]. One of us has recently shown that active noise also enhances particle diffusion in the crowded cytoplasm, and this enhancement effect is particularly significant for large particles [35]. Here, we find that particle diffusion can still be suppressed in the presence of active noise in cytoplasm far above jamming (Figure S6), which can be another reason that real cytoplasm is not far above jamming.

One should note that in this work, we study the viscoelasticity of cytoplasm and compare our data with microrheology experiments in which the cytoskeleton does not affect the probe particle [11, 13]. It has been shown that the cytoskeleton can reduce the exponent between the shear modulus and frequency from 0.5 to 0.1 [15], consistent with measurements involving the cytoskeleton [3, 8, 36]. We note that in Ref. [15], the authors proposed an alternative model using a self-similar network of springs and dashpots to explain the origin of the *ω*^1*/*2^ scaling of the complex modulus, distinct from the mechanism based on the jamming transition of colloidal suspension. We remark that besides rationalizing the *ω*^1*/*2^ scaling of complex modulus, our colloidal model also predicts that a finite active noise makes the cytoplasmic viscosity less sensitive to changes in volume fraction, in agreement with experiments of live cells [8, 10].

Experiments based on osmotic perturbations have estimated the volume fraction of osmotically inactive components to be about 30% [37]. We argue that this may not conflict with our conclusions because the repulsive interaction between proteins is set by their gyration radii [38]. Considering the widely existing intrinsically disordered regions across the proteome, this length scale can be much longer than the expectation by considering proteins as compact objects [39].

Are there any other advantages for cells to be in the critical state besides being sensitive to active noise? One should note that the storage and loss modulus are approximately equal over a wide range of frequencies since they share the same scaling with the frequency. Therefore, the phase shift between stress and strain maintains *π/*4 over a wide range of frequencies, as confirmed numerically (Figure S7, S8). This robustness may give cells an additional evolutionary advantage. Theoretically, our conclusions in this work are equally valid in two-dimensional systems as expected (Figure S9-S13) [40]. Finally, tuning active noise is not the only way to change the cytoplasmic viscosity. For example, Ref.[41] showed that budding yeast can modulate viscosity by regulating the synthesis of glycogen and trehalose. The variety of regulation mechanisms suggests the complexity behind viscosity adjustment and requires further investigation.

## Acknowledgements

We thank Lingyu Meng, Yiyang Ye, and Hua Tong for helpful discussions about this work. The research was supported by Peking-Tsinghua Center for Life Sciences grants.

