## Supplemental Material for "Ultrasensitive tuning of cytoplasmic viscosity via active noise"

### Supplementary Material

Ziming Zhao[1] and Jie Lin[1,2,3]
[1]School of Physics, Peking University, Beijing, China
[2]Center for Quantitative Biology, Peking University, Beijing, China
[3]Peking-Tsinghua Center for Life Sciences, Peking University, Beijing, China
(Dated: December 5, 2024)

#### NUMERICAL SIMULATION DETAILS

We perform numerical simulations in a cubic space by solving the overdamped Langevin equation of each particle with the periodic boundary condition in C++. Simulations for a single 3D packing are in an $80 \times 80 \times 60$ region, and simulations for a single 2D packing are in a $400 \times 400$ region, both involving $\sim 3 \times 10^4$ particles. To avoid crystallization, a 50-50 mixture of particles with a size ratio of 1.4 are initially randomly placed in space [1]. Before we apply a shear, we first let the system evolve for $t = 5 \times 10^3$ to reach a steady state [2]. After removing any residual strain to stabilize the system [3], we apply a small strain $\epsilon = 10^{-2}$ to the system through the Lees-Edwards boundary condition. Both in 2D and 3D experiment, we take a time step $\Delta t = 10^{-3}$ until $T = 10^6$, and record the stress $\sigma(t)$. To ensure the stability of the results, at least 32 independent simulations with random initial conditions were performed under each situation.

In particle-diffusion simulations (S6), we assume that the amplitude of the active noise is proportional to the cube of nondimensional particle radius $a_i$:

$$\langle \xi_{i,\alpha}(t)\xi_{j,\beta}(t')\rangle = 2\delta_{ij}\delta_{\alpha\beta}\eta_i k_B(T_{\mathrm{p}} + T_{\mathrm{a}}a_i^3)\delta(t-t'). \tag{S1}$$

This form of active noise is consistent with experimental data of *E. coli* [4]. One of us has shown that a random force with a constant magnitude whose pointing direction diffuses due to thermal fluctuation has an autocorrelation function proportional to the cube of particle radius [5].

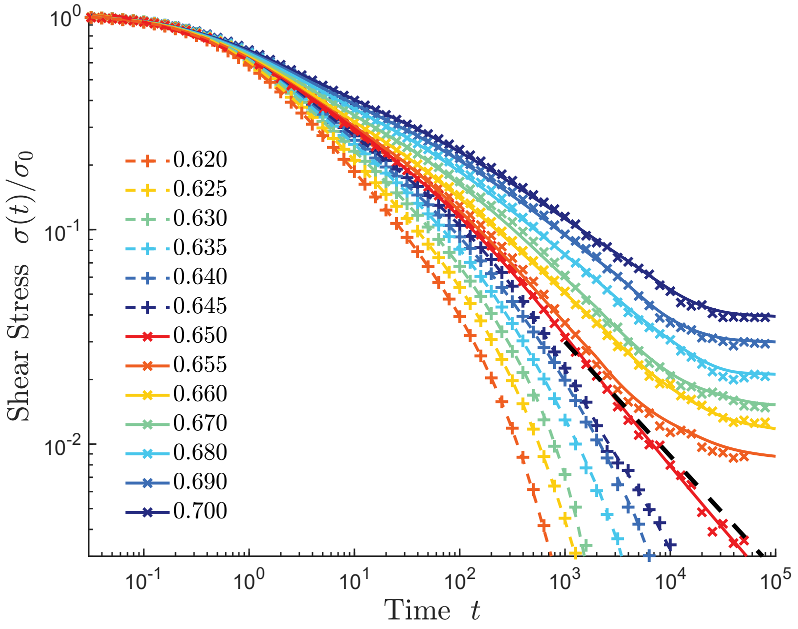


FIG. S1. The relaxation curve with different volume fractions without any noise, $T_{\text{eff}} = 0$. Below the jamming volume fraction $\phi_J \approx 0.650$, the shear stress relaxes to zero, while for systems above jamming, the stress approaches a finite residual stress. At the jamming point $\phi = \phi_J$, the stress decays as $\sigma(t) \sim t^{-1/2}$. The black dashed line has a slope of $-0.5$ in the log-log plot. The colorful dashed lines represent a third-order polynomial fitting of the simulated data. $\sigma_0$ is the shear stress at time 0.

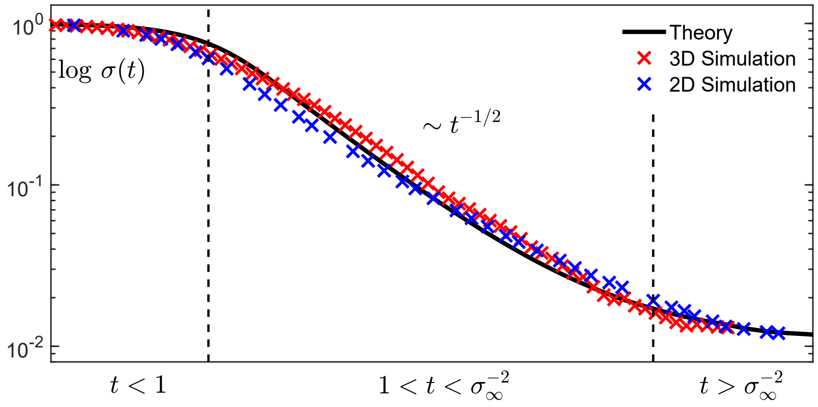


FIG. S2. Comparison between the simulated athermal relaxation curves in 3D (red cross) and 2D (blue cross) and the numerical calculation of Eq.(4) in the maintext with a finite residual stress. The formula Eq.(4) catches the the property of platform $\sigma(t) \approx 1$ when $t < 1$, $\sigma(t) \sim t^{-1/2}$ when relaxing and finite residual stress $\sigma \to \sigma_\infty$ in the long time limit. Here, we choose volume fraction $\phi_{2\text{D}} \approx 0.844$ and $\phi_{3\text{D}} \approx 0.675$ for 2D and 3D simulations, respectively and they have similar $\sigma_\infty \approx 0.011$.

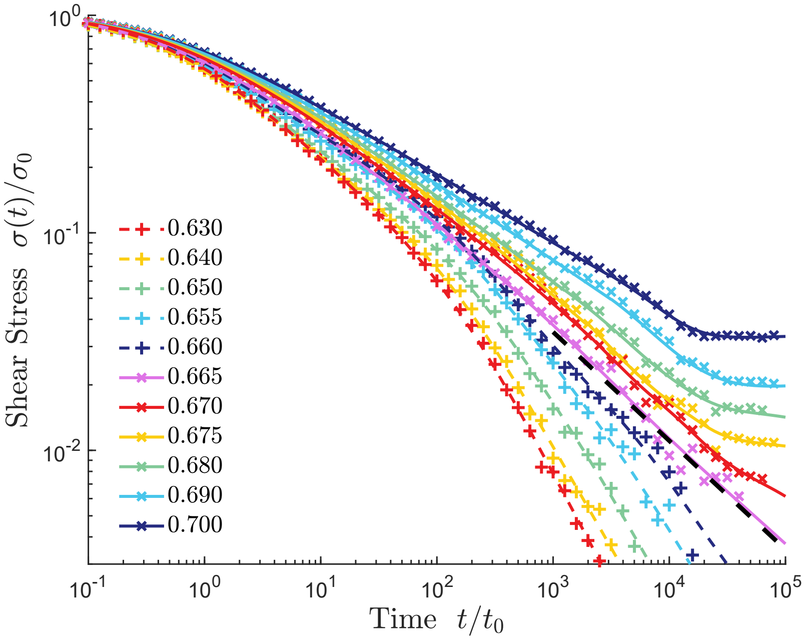


FIG. S3. The relaxation curve with different volume fractions $\phi$ at $kT_{\mathrm{eff}} = 1.8 \times 10^{-4}$. At $\phi = 0.665$, the stress decays as $\sigma(t) \sim t^{-1/2}$. The black dashed line has a slope of $-0.5$ in the log-log plot.

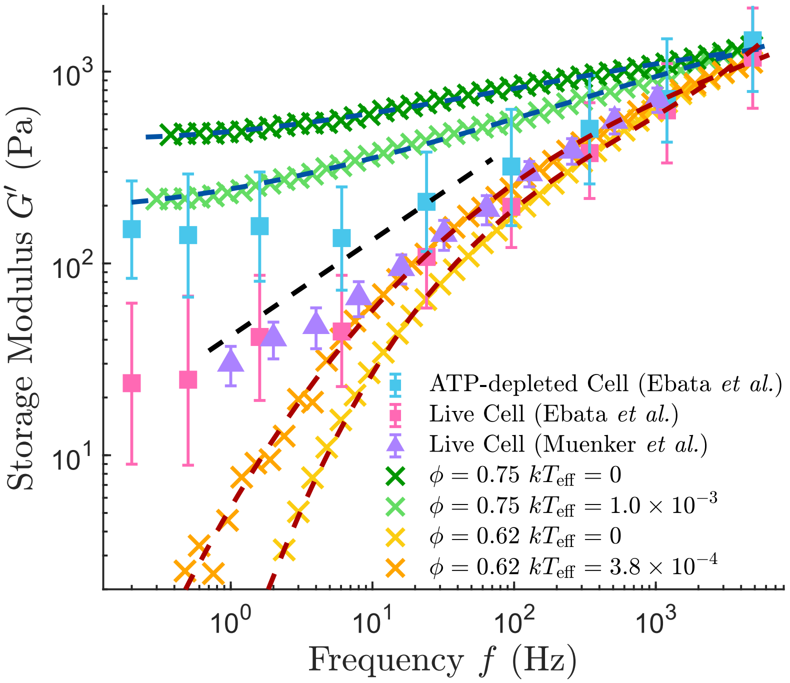


FIG. S4. We compared the simulated storage modulus of $\phi = 0.750$, $kT_{\mathrm{eff}} = 10^{-3}$ and $\phi = 0.620$, $kT_{\mathrm{eff}} = 0$ with experiment data and finding a significantly different increase slope. According to our results, active force must be nearly comparable with elastic energy $\kappa a_0^2$ to sufficiently fluidized a highly jammed system, which is unreasonable in a biological context.

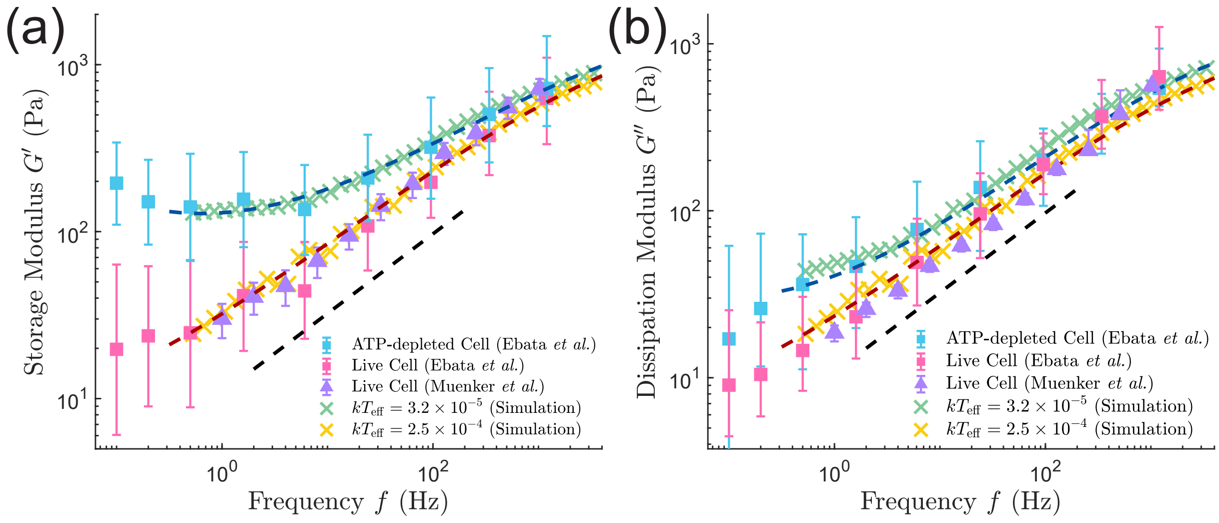


FIG. S5. The simulated complex moduli with $\phi = 0.675$ at the critical effective temperature $T_{\rm eff,c}$ match the experimental data well, including the storage modulus (a) and the loss modulus (b). The active part of the effective temperature is again dominant over the passive part.

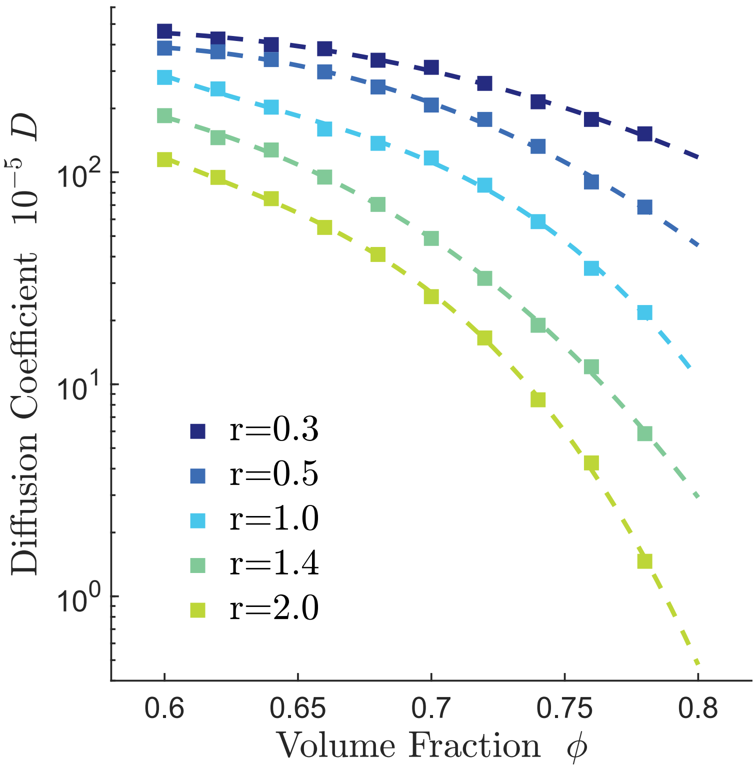


FIG. S6. Diffusion coefficient for particles with different radius, under both passive and active force as volume fraction increases. We take active force and passive force as $kT_{\rm a} = 10^{-3}$ and $kT_{\rm p} = 10^{-4}$ respectively in simulations with a 50-50 mixture of particles with size ratio of 1.4 as background and test particle diffusion. We find that as volume fraction grows above jamming, transportation for large particles are more strictly suppressed than small particles. The diffusion enhancement driven by active energy cannot compensate for the suppression caused by a highly jammed volume fraction. Therefore, considering diffusion efficacy, volume fraction cannot grow limitlessly.

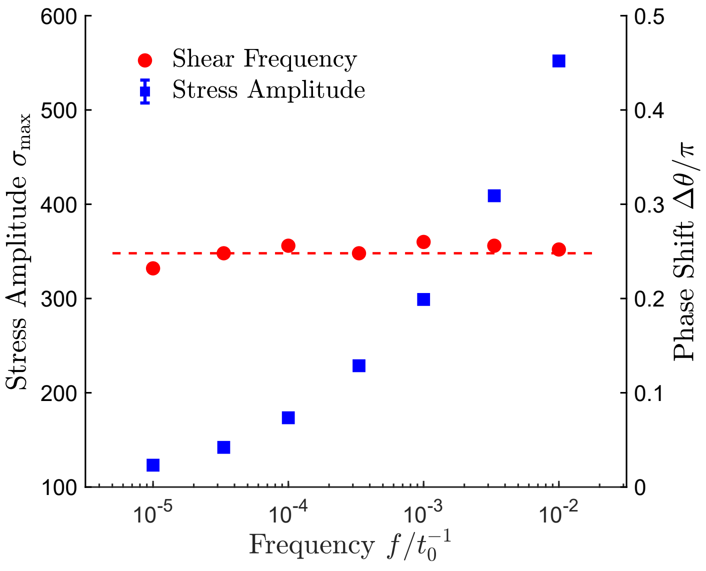


FIG. S7. Relative amplitude (blue) and phase shift (red) of stress under strain of different frequencies. As predicted, we observe an invariant phase shift $\Delta\theta = \pi/4$, which may provide a potential evolutionary advantage. A sinusoidal shear $\epsilon = \epsilon_0 \sin(2\pi f t)$ is applied with frequency $f$ and period $T_0 = f^{-1}$. The stress becomes $\sigma(t) = \sigma_{\max}(2\pi f t + \Delta\theta)$ with an amplitude $\sigma_{\max}$ and phase shift $\Delta\theta$.

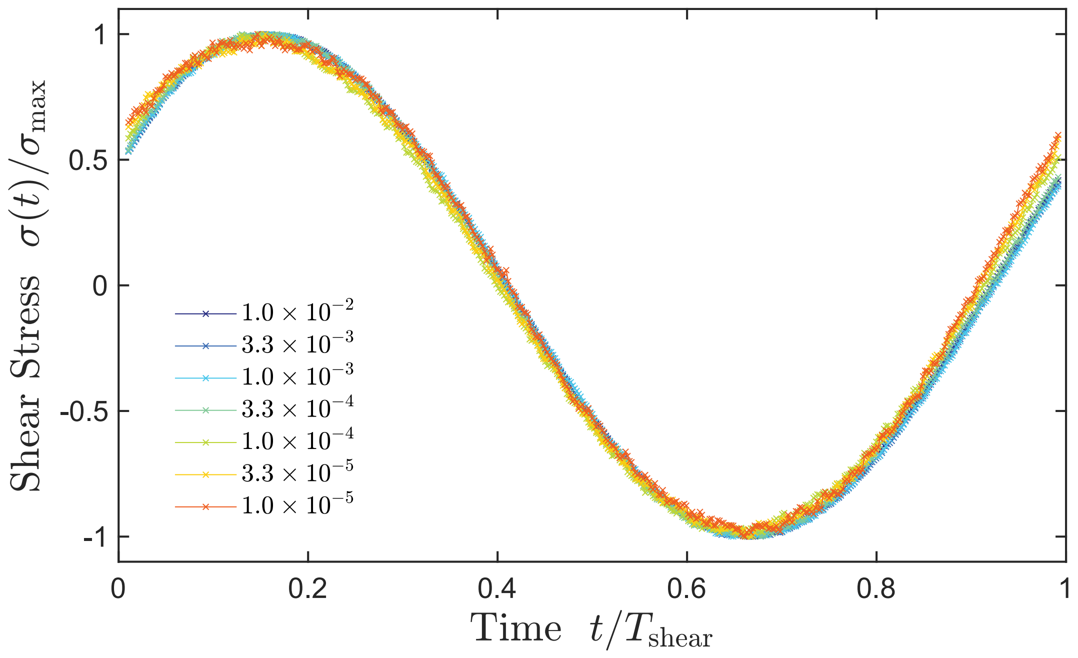


FIG. S8. Stress vs. time under periodic strain. The vertical and horizontal axis are normalized with the period $T_0$ and the maximal stress $\sigma_{\max}$. We observe identical phase shifts independent of strain frequency.

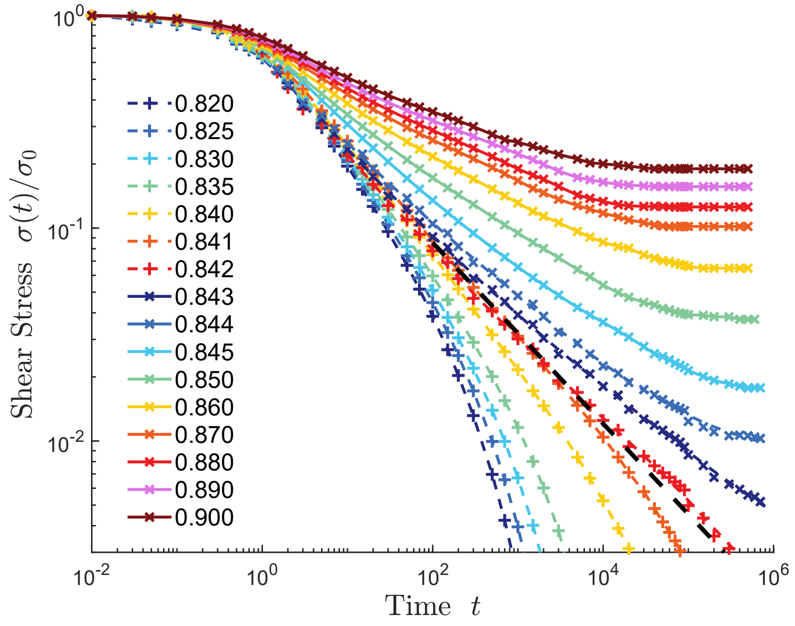


FIG. S9. The relaxation curve with different volume fractions without any noise, $T_{\rm eff} = 0$ in 2D. Below the jamming volume fraction $\phi_J \approx 0.842$, the shear stress relaxes to zero, while for systems above jamming, the stress approaches a finite residual stress. At the jamming point $\phi = \phi_J$, the stress decays as $\sigma(t) \sim t^{-1/2}$. The black dashed line has a slope of $-0.5$ in the log-log plot. The colorful dashed lines represent a third-order polynomial fitting of the simulated data. $\sigma_0$ is the shear stress at time 0.

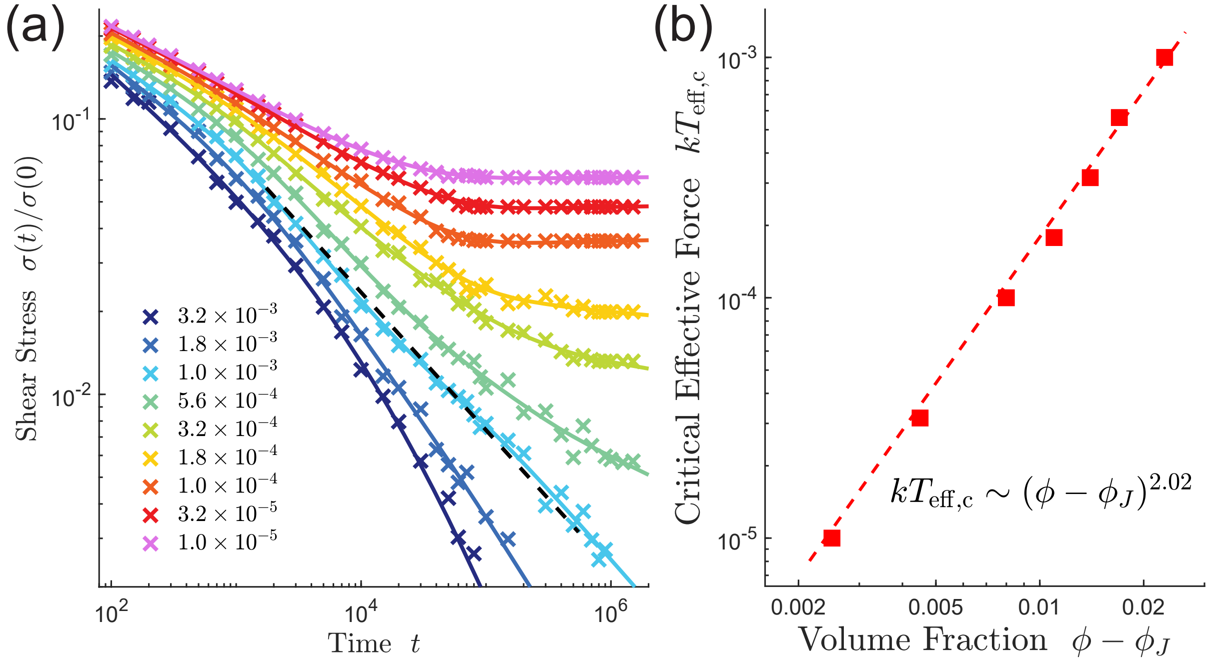


FIG. S10. (a) A jammed cytoplasm with $\phi = 0.860$ in 2D is fluidized by noise such that the stress relaxes to zero at a high enough effective temperature. The critical effective temperature $T_{\rm eff,c} \approx 1.0 \times 10^{-3}$ in this case. The colorful dashed lines represent a third-order polynomial fitting of the simulated data. The black dashed line has a slope of $-0.5$ in this log-log plot. (b) The scaling between the critical temperature $T_{\rm eff,c}$ and $\phi - \phi_J$ where $\phi_J$ is the volume fraction at the jamming transition at zero temperature.

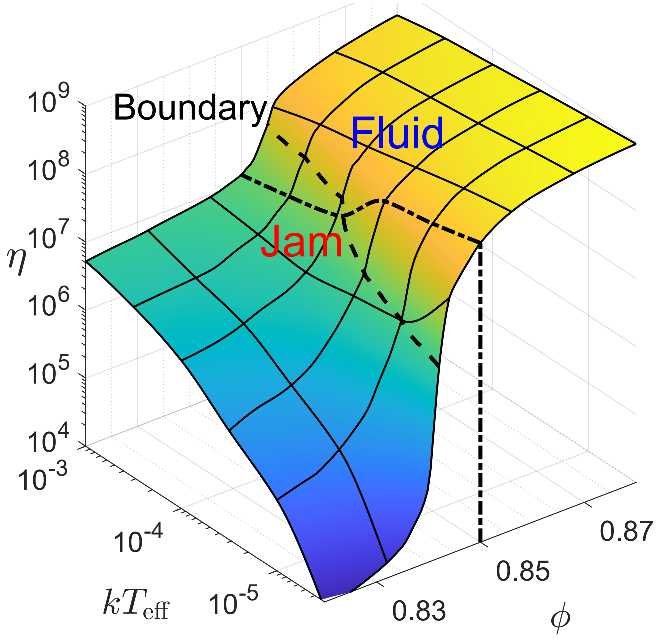


FIG. S11. Viscosity as a function of the volume fraction and effective temperature in 2D. The dot dash line represents the viscosity for $\phi = 0.85$. The dashed line represents the transition boundary for critical temperature $T_{\rm eff,c}$.

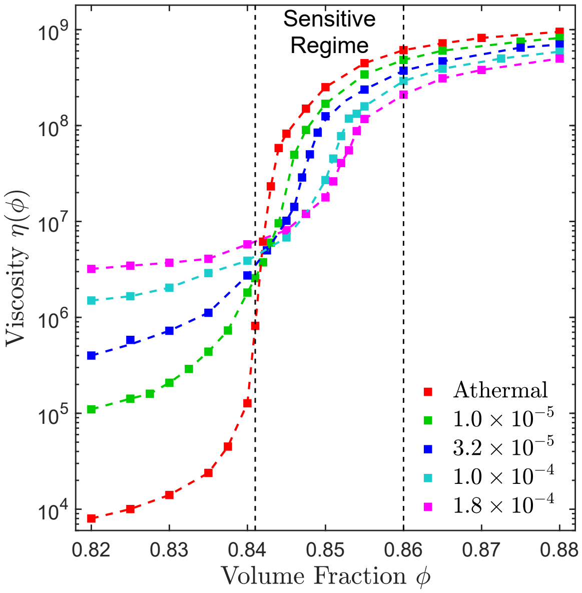


FIG. S12. Viscosity vs. $\phi$ under different effective temperatures in 2D. We highlight the sensitive regime slightly above the jamming point $\phi_J \approx 0.842$. In this regime, a finite active noise also makes the viscosity less sensitive to the change in $\phi$.

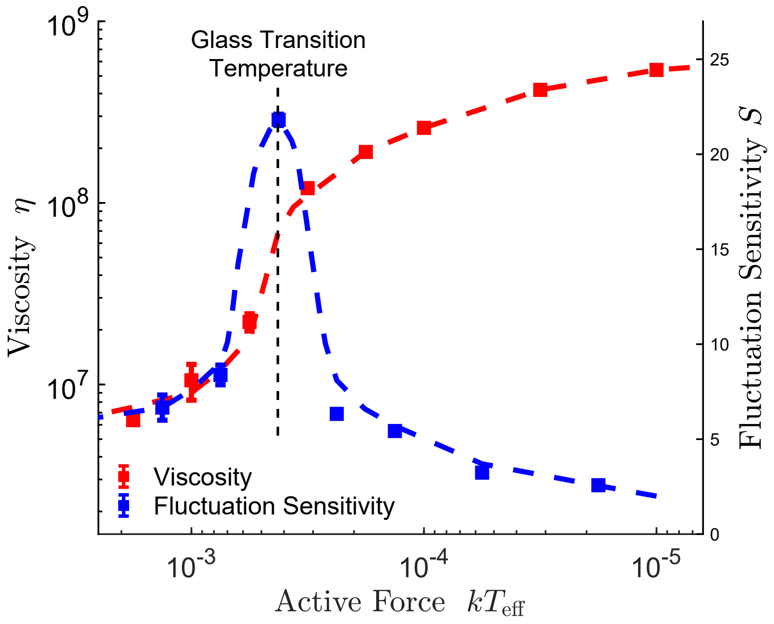


FIG. S13. Viscosity vs. $T_{\mathrm{eff}}$ for $\phi = 0.850$ on a log-log plot for 2D situations. Sensitivity $S$ is defined as the ratio between the rate of temperature change and the rate of viscosity change. Note that sensitivity peaks at $kT_{\mathrm{eff,c}} \approx 4 \times 10^{-4}$, which is the critical effective temperature for the system.
